# Transmission delays and frequency detuning can regulate information flow between brain regions

**DOI:** 10.1101/2020.07.09.194969

**Authors:** Aref Pariz, Ingo Fischer, Alireza Valizadeh, Claudio Mirasso

## Abstract

Brain networks exhibit very variable and dynamical functional connectivity and flexible configurations of information exchange despite their overall fixed structure (connectome). Brain oscillations are hypothesized to underlie time-dependent functional connectivity by periodically changing the excitability of neural populations. In this paper, we investigate the role that the connection delay and the frequency detuning between different neural populations play in the transmission of signals. Based on numerical simulations and analytical arguments, we show that the amount of information transfer between two oscillating neural populations can be determined solely by their connection delay and the mismatch in their oscillation frequencies. Our results highlight the role of the collective phase response curve of the oscillating neural populations for the efficacy of signal transmission and the quality of the information transfer in brain networks.

**Author summary:** Collective dynamics in brain networks is characterized by a coordinated activity of their constituent neurons that lead to brain oscillations. Many evidences highlight the role that brain oscillations play in signal transmission, the control of the effective communication between brain areas and the integration of information processed by different specialized regions. Oscillations periodically modulate the excitability of neurons and determine the response those areas receiving the signals. Based on the communication trough coherence (CTC) theory, the adjustment of the phase difference between local oscillations of connected areas can specify the timing of exchanged signals and therefore, the efficacy of the communication channels. An important factor is the delay in the transmission of signals from one region to another that affects the phase difference and timing, and consequently the impact of the signals. Despite this delay plays an essential role in CTC theory, its role has been mostly overlooked in previous studies. In this manuscript, we concentrate on the role that the connection delay and the oscillation frequency of the populations play in the signal transmission, and consequently in the effective connectivity, between two brain areas. Through extensive numerical simulations, as well as analytical results with reduced models, we show that these parameters have two essential impacts on the effective connectivity of the neural networks: First, that the populations advancing in phase to others do not necessarily play the role of the information source; and second, that the amount and direction of information transfer dependents on the oscillation frequency of the populations.

## Introduction

A typical sensory response process in the nervous system consists of the active selection of relevant inputs, the segregation of the different features of the input, and the integration of the information leading to the right action. All these stages depend on the flexibility in the information routing, as well as in an efficient communication between different regions of the nervous system. However, the circuit and the dynamical mechanism explaining the fast reconfiguration of the effective pattern of communication and the information transfer in the neural systems have so far not been satisfactorily understood.

One interesting and widespread proposal is that in the presence of neural oscillations, communication patterns can be controlled by adjusting the phase relation between the local oscillations of different brain regions [1–8]. In the brain, synaptic interactions lead to correlated activity of the neurons and the appearance of successive epochs of high and low excitability, characterized by collective neuronal oscillations in different frequency bands [9–15].

Neural oscillations establish intermittent windows of high and low excitability, giving rise to a time-dependent response of the system to the inputs from other brain regions. According to the communication through coherence (CTC) hypothesis, it is possible to adjust the phase relationship between two regions to activate and deactivate the communication channel [1, 16–18] or continuously vary the efficacy of the channel [5]. While in the original proposal, the widespread variability in neural systems and the inconsistency of the coherence across time and space were ignored, recent studies showed that the mechanism works if oscillations are not persistent and even if the locking is not stable [19, 20].

The diversity and the time-dependency of the phase relationships reported in experiments [5, 21–23] are supposed to underlie the rich variety of communication patterns in the nervous circuits. Several experimental and computational studies have shown that those regions that phase advance others act as leaders and can efficiently transmit information to the laggard regions [20, 24, 25]. It has been shown that the presence of frequency mismatch can give rise to a phase difference and a directional information transfer, that is, nodes with higher frequency transmit information to those with lower frequency when the connection delay is neglected [20, 24]. However, one of the key parameters which determine synchronization and the phase relation between coupled oscillators, is the interaction delay due to the finite time of transmission of signals between the oscillators. Since the synchronization and the phase relations determine the effective routes for information transfer, and the synchrony properties depend on the interaction delays, an important question arises: How do transmission delays in brain circuits affect the effective communication patterns?

In many theoretical and computational studies in networks of coupled dynamical systems, delays are disregarded mainly due to the analytical complexity and computational burden. However, delays might have a crucial impact on the collective properties of distributed dynamical systems [6, 7, 26–30]. In the brain, delays in the transmission of signals between neurons and neural populations are quite heterogeneous and cover a wide range, from milliseconds to tens of milliseconds [31]. So they can be of the same order or larger than some important neural time scales, for example, the integration time of the membrane, the period of gamma oscillations, or even of other bands, and temporal window for spike timing dependent plasticity [13, 32–38]; therefore cannot be ignored. Consequently, an adequate treatment of the CTC theory, needs the delays to be explicitly taken into account. In fact, the phase relation resulting in maximum signal transmission efficiency is not a zero phase-lag when the transmission delays are non-negligible [39, 40].

In this manuscript, we study the conditions for an effective communication between two coupled neural populations by systemically varying the interaction delay and mismatch of their oscillation frequencies. Our results show that for small delays, information encoded in the population oscillating at higher frequency is transmitted to other populations and when the information is encoded in the population oscillating at lower frequency, the other population is unable to receive the information; compatible with previous results [24, 41]. We find, however, that this is not always the case and the amount and direction of the effective communication between populations depend, in general, on the interaction delay. Symmetric information transmission and efficient transmission in reverse direction (from slow to fast) are also passible for certain delays.

Moreover, a formulation based on the coupled phase oscillators and the phase response curve allows us to provide a general framework to predict how the pattern for effective communication between two coupled oscillators, changes with delay and frequency mismatch. This novel findings provides a theoretical basis for how the information is transmitted in the brain circuits along different channels and directions and over different frequency bands.

## Materials and methods

### Neuron model

In our simulation we used the Hodgkin-Huxley (HH) neuron model [42]. The evolution of the membrane potential and gate variables are given by:

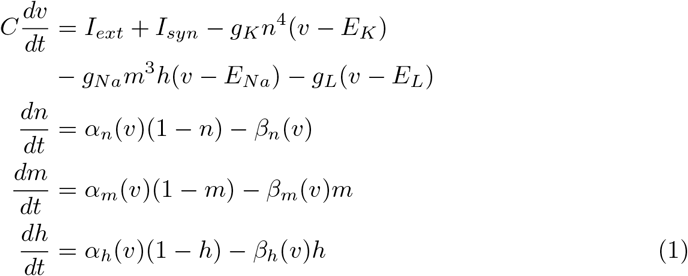

The *α_x_* and *β_x_, x* ∈ (*n, m* and *h*) are defined as below

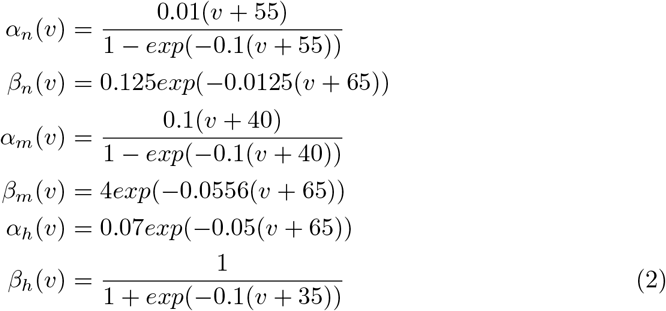

where *I_ext_* and *I_syn_* are the input and synaptic currents, respectively. The values of parameters are given in Table 1.

**Table 1.** Simulation parameters.

| Parameter | Value | Description |
| --- | --- | --- |
| C | 1 $\mu F/cm^2$ | Capacitance |
| $g_K$ | 36 $mS/cm^2$ | K conductance |
| $g_{Na}$ | 120 $mS/cm^2$ | Na conductance |
| $g_L$ | 0.3 $mS/cm^2$ | Leak conductance |
| $E_K$ | -77 *mV* | K reversal potential |
| $E_{Na}$ | 50 *mV* | Na reversal potential |
| $E_L$ | -54.4 *mV* | Leakage reversal potential |
| $E_{syn}^E$ | 0 *mV* | Excitatory reversal potential |
| $E_{syn}^I$ | -80 *mV* | Inhibitory reversal potential |
| $t_d^{inner}$ | 0 – 14 *ms* | Inner population delay of excitatory unit |
| $t_d^{intra}$ | 0.5 *ms* | Intra population axonal delay |
| $\tau_d$ | 3 *ms* | Synaptic decay time |
| $\tau_r$ | 0.5 *ms* | Synaptic rise time |
| $I_{ext}$ | 10 – 12 $\mu A/cm^2$ | Injected current |
| $\bar{\mu}, \sigma$ | 0, 0.5 $\mu A/cm^2$ | Mean and variance of Gaussian white noise |
| $g_{EE}$ | 3.75 $\mu S/cm^2$ | Synaptic weight, E → E |
| $g_{EI}$ | 15 $\mu S/cm^2$ | Synaptic weight, I → E |
| $g_{IE}$ | 7.5 $\mu S/cm^2$ | Synaptic weight E → I |
| $g_{II}$ | 15 $\mu S/cm^2$ | Synaptic weight I → I |

The synaptic current of the i-th neuron 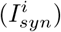 is given by:

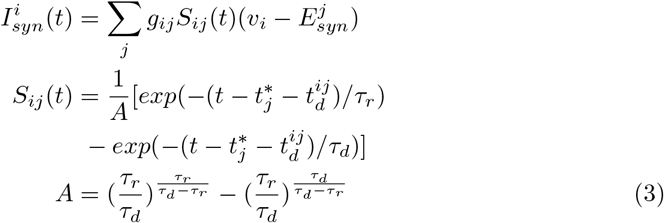

*v_i_* is the membrane potential of the post-synaptic neuron and 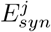 is its reversal synaptic potential. *S_ij_* is a double-exponential function, modeling the efficacy of the chemical synapses mediated by *AMPA* and *GABA_A_* receptors. 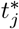 is the time at which the pre-synaptic neuron spikes and *t_d_* is the axonal delay between pre- and post-synaptic neurons. The synapses’ parameters and synaptic weights (*g_ij_*) are given in Table 1.

We numerically solved the equations using the Milshtein algorithm with an integration step *dt* = 0.01 ms.

### Population architecture

Each of our populations was composed of 100 neurons with 80% excitatory and 20% inhibitory neurons. The connectivity within the population was chose as random and with probability of 10%. The connectivity between populations was also chosen as random (but just among excitatory neurons) with probability 5%. The intra population delay was taken as 0.5 *ms* while that between populations was varied from 0 to 14 *ms*.

### Input signals

We inject constant current (varied from 10 to 12 *μA/cm*^2^) and a Gaussian white noise (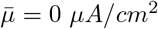, *σ* = 0.5 *μA/cm*^2^) to each neuron (*I_ext_* in Eq. 1). Due to the value of the input current, neurons fired within a frequency range of 70-73 Hz.

To test the signal transmission we injected slow (5 Hz) and fast (70 Hz) non-periodic signals (Depicted in Fig. 7) into the excitatory neurons of the host (or sender) population. By running simulations in the absence of an external signal, we found the maximum firing rates. Its inverse value characterizes the oscillation period. In the fast signal case, we applied a single pulse at a certain phase in a period, on all the excitatory neurons. The phase of the impact of the pulse changed in each experiment by dividing the oscillation cycle into 50 segments. In each experiment, we applied a rectangular pulse of amplitude *I_pulse_* = 0.25*μA/cm*^2^ and width of 2 (ms).

**Fig 1.**
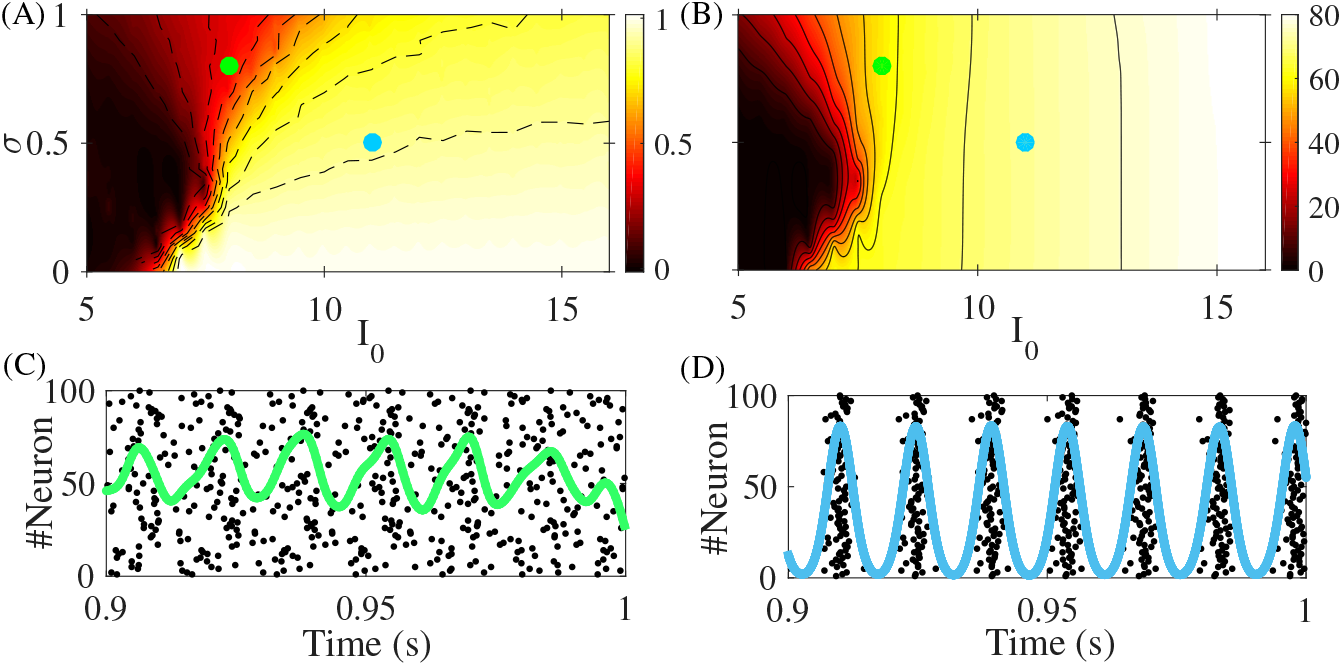
Properties of a single population. The coherency (panel (A)) and the oscillation frequency (panel (B)) of the population activity is plotted in color code in the noise amplitude (*σ*) vs. mean input current (*I*_0_) plane. In panel C (D) the raster plot and the activity of a single population for *I*_0_ = 8 (*I*_0_ = 11) and *σ* = 0.8 (*σ* = 0.5) is plotted with green (blue) line. In the rest of the simulations we use the set of parameters considered in panel (D) resulting in a coherency value of 0.80 (green point in panel (A)).

**Fig 2.**
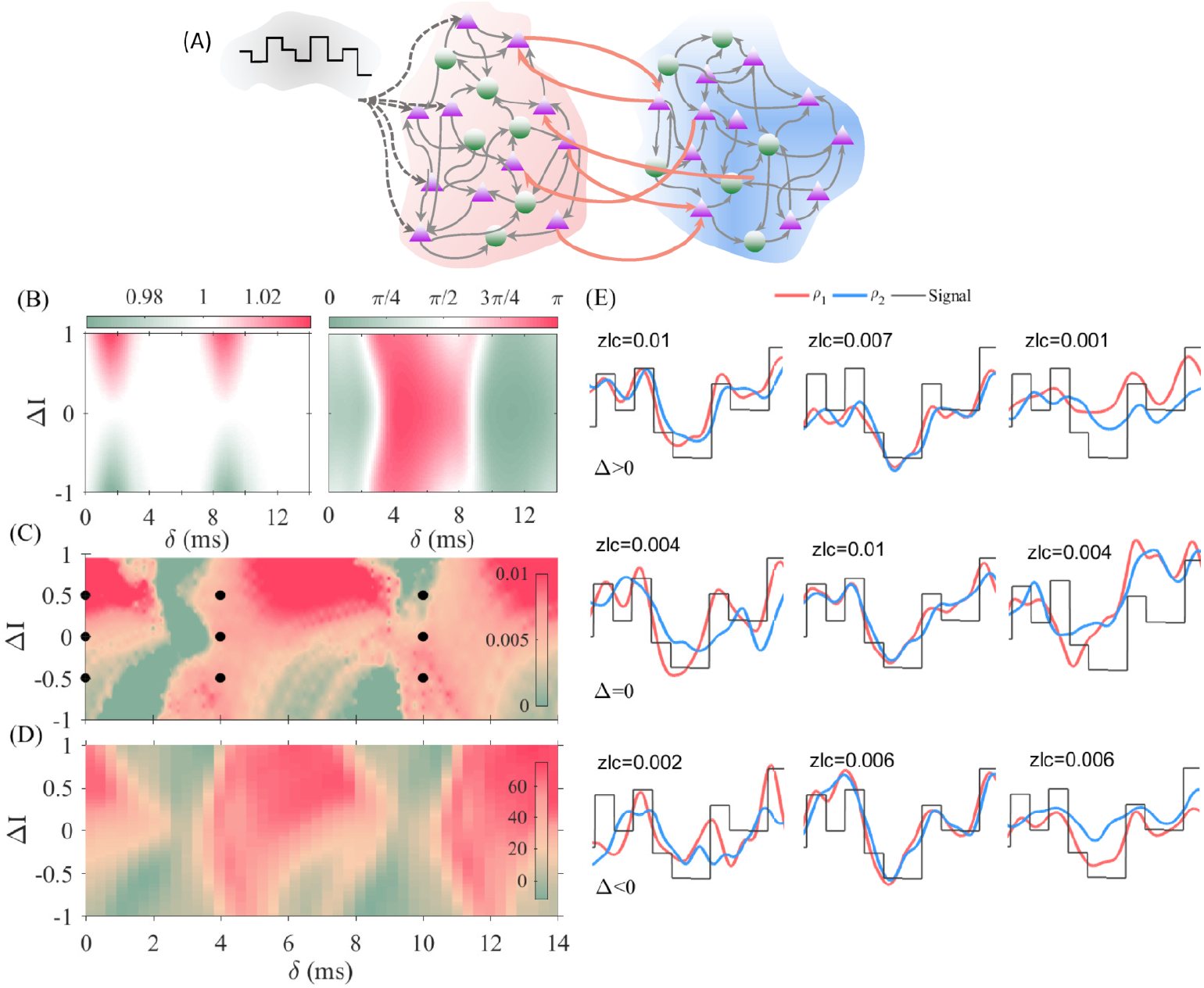
Signal transmission and information transfer between the two bidirectionally coupled populations. (A) Schematic diagram of the network connectivity. Only excitatory neurons between the two populations are connected. (B) The locking zone (left panel) and phase difference (right panel) of the activity of the two populations are shown in color code in the input current mismatch (Δ*I*) vs. delay (δ) plane. Zero-lag cross-covariance (ZLC) between the firing rate of the second population with the input signal (C), and the delayed mutual information between firing rates of two populations (D) are shown in color code in the same plane as in panel (B). In (E) the firing rate of the two populations and the input signal are plotted, for the parameters values marked with black dots in (C). The value of ZLC is shown in each panel. The offset and the amplitude of the external signal were scaled for a better comparison.

**Fig 3.**
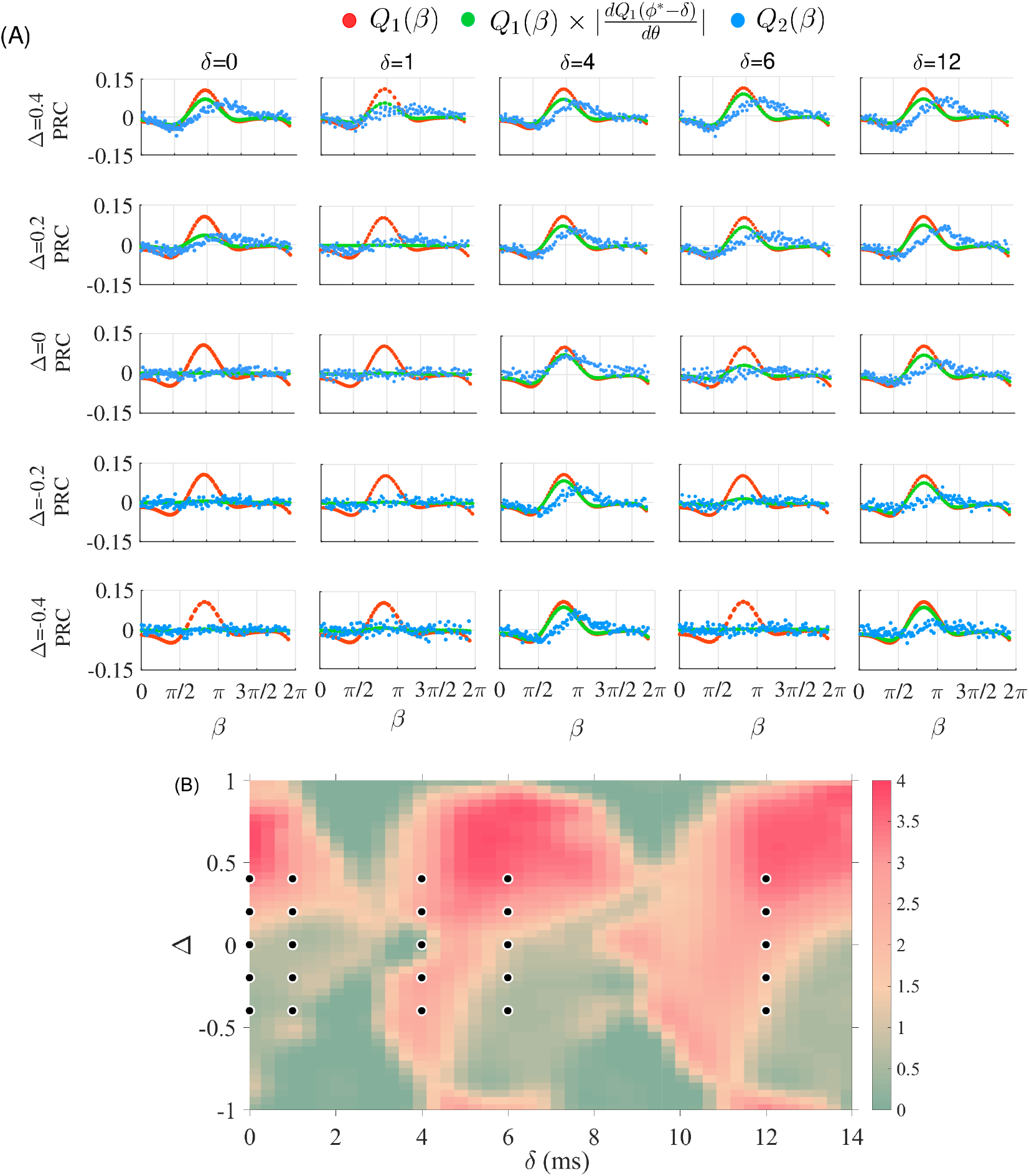
Response of the network to fast signals. (A) The PRC of the sender population (pPRC; red) and the PRC of the receiver population (non-local PRC, nPRC; sky-blue) is plotted versus the phase *β* for different values of the detuning Δ. The green curve shows the analytical prediction for the response calculated as the value of the derivative of PRC, multiplied by the value of PRC. To compute the nPRC we use the phase (or time) at which the pulse is injected in the sender population, and calculate the difference of the ISIs without and with signal injection. (B) The integral of the absolute value of the response of the receiver population to the input pulse on the sender population (area under the absolute value of blue curves in (A)) is ploted in the detuninng vs. delay plane.

**Fig 4.**
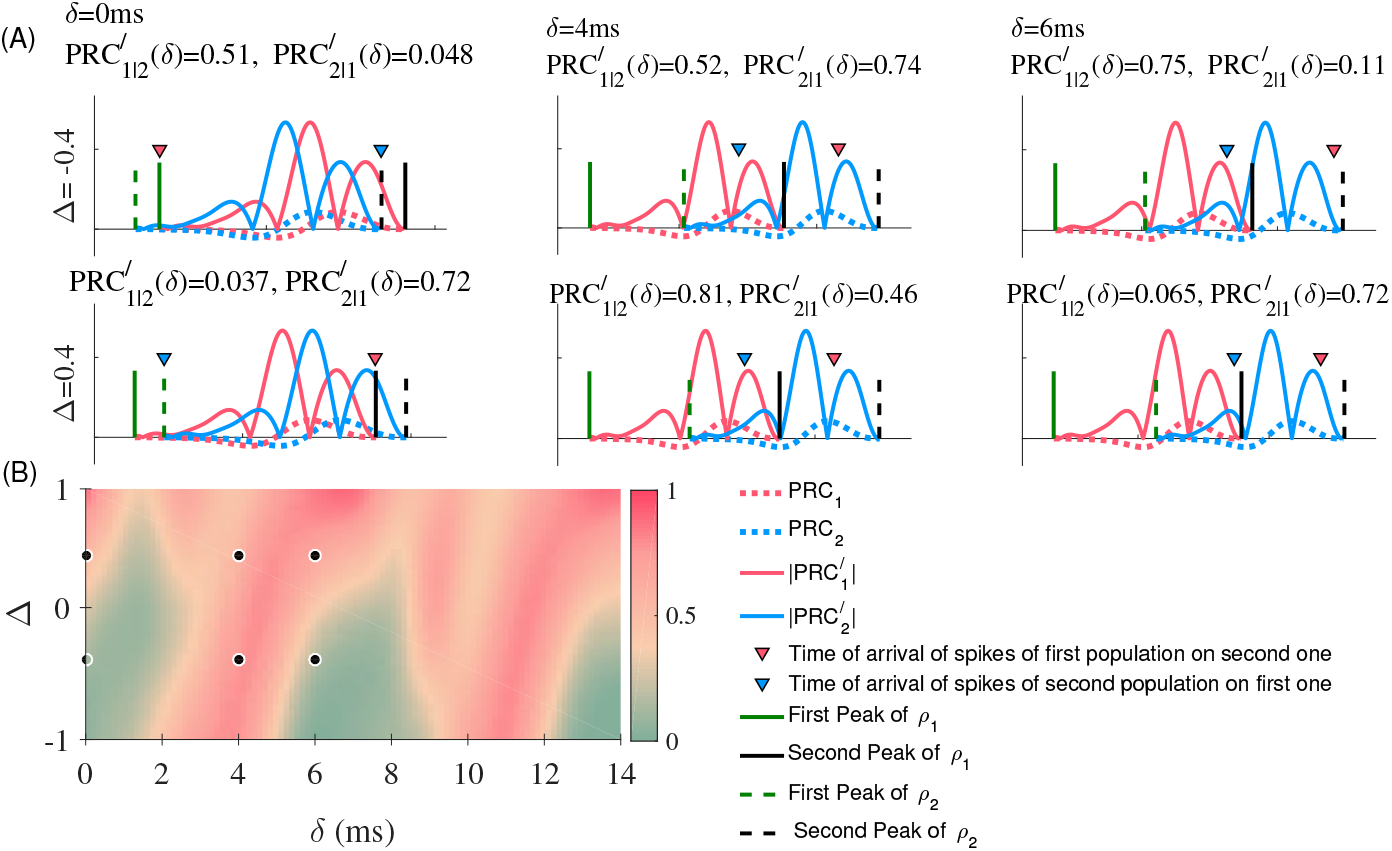
(A) Schematic plot of the PRC (dotted lines) and its time derivative (solid lines) represented between two successive oscillation peaks of the populations for three delays, *δ* = 0, 4, 6 (ms) and two detuning values, Δ = –0.4, 0.4. The values on top of each panel 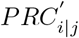, are the absolute value of *PRC*′ of the population *i* when the spikes of population *j* arrives. This measure predicts the quality of the transmission. (B) The absolute value of derivative of the PRC of the receiver population at the time it receives the spikes of the sender population, is plotted in the detunning vs. delay plane.

**Fig 5.**
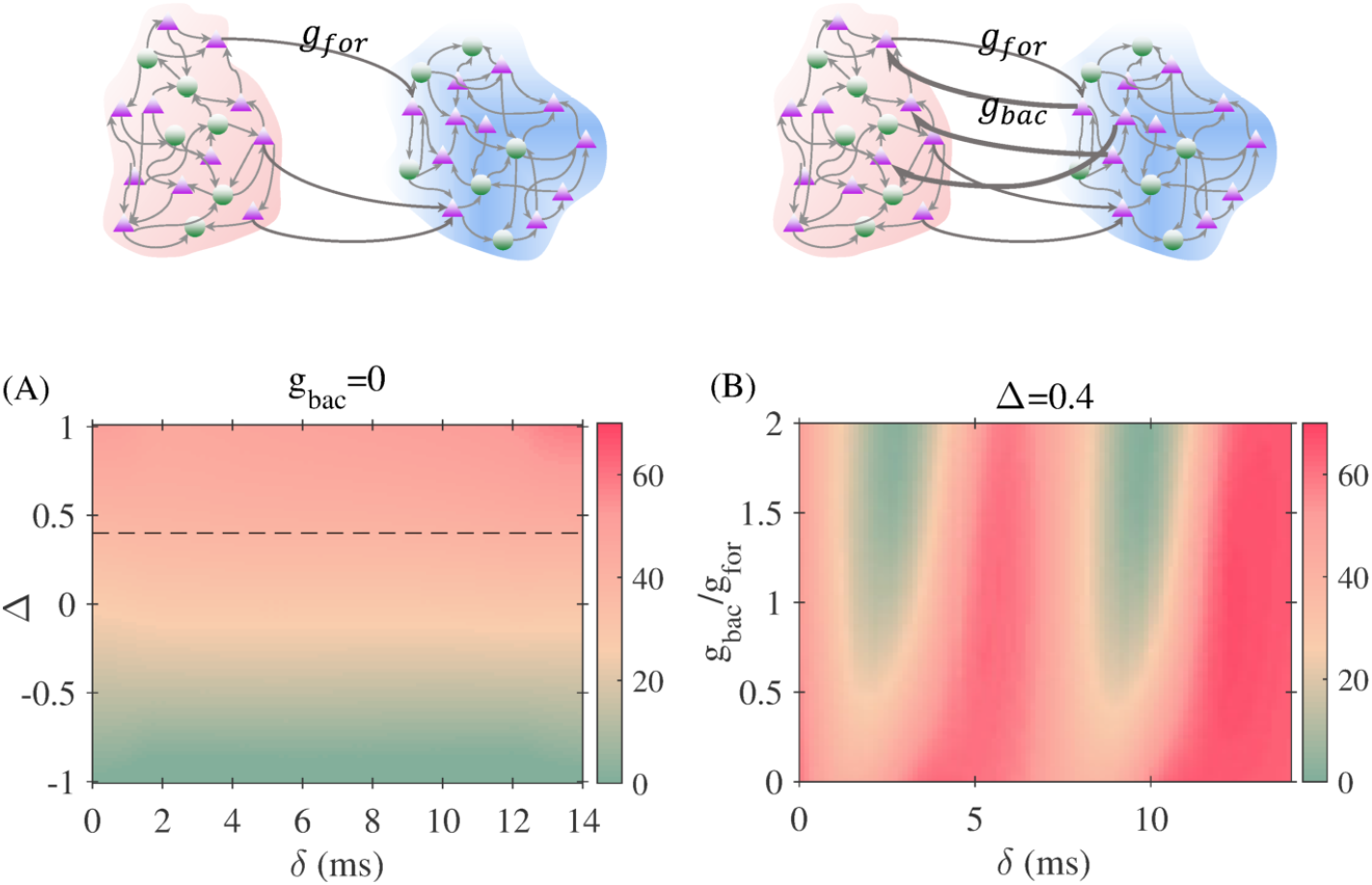
Effect of connection asymmetry on the information transfer. In (A) the connection is unidirectional from the sender to the receiver. The mutual information transfer depends on the detunning but, as expected, is independent of the delay. In (B) the delayed mutual information is plotted, in color code, in the ratio between feedforward and feedback strength versus delay plane for a positive inhomogeneity value (Δ= 0.4).

**Fig 6.**
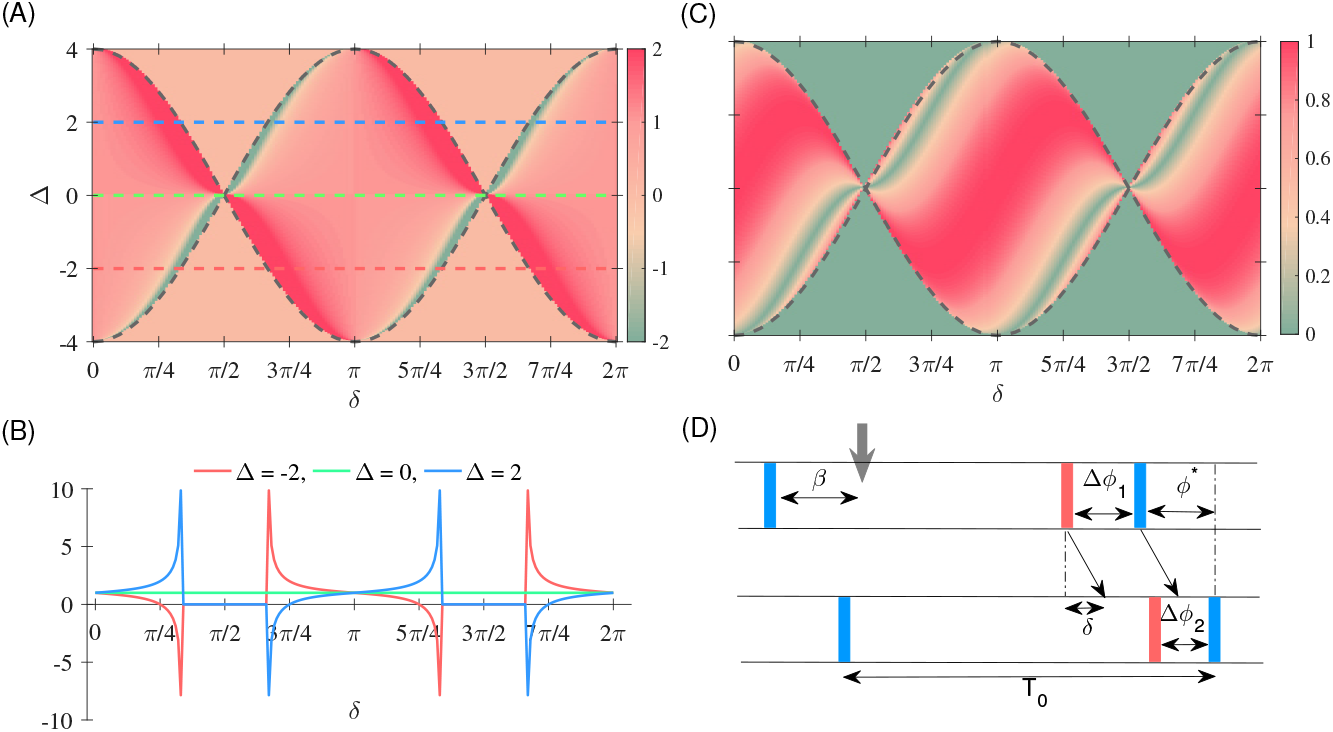
Analytical results. (A) The response function *S_i_* (Eq. 11) to the input of slow signals for the case of two delay-coupled phase oscillators is plotted in color code, in the detuning vs. phase plane. In (B) the same results as (A) are shown for Δ = –2,0, 2, depicted by red, green and blue colors in (A), for the sake of clarification. (C) The absolute value of derivative of the PRC (Q), at phase (*ϕ*^*^ – *δ*). To plot this figure, we first find for each value of Δ and *δ* the locked regime using the Eq. 9 and then we find the absolute value of sin(*ϕ*^*^ – *δ*) (D) The schematic of how a pulsatile perturbation in the first oscillator affects the phase of the second oscillator, as discussed in the text.

**Fig 7.**
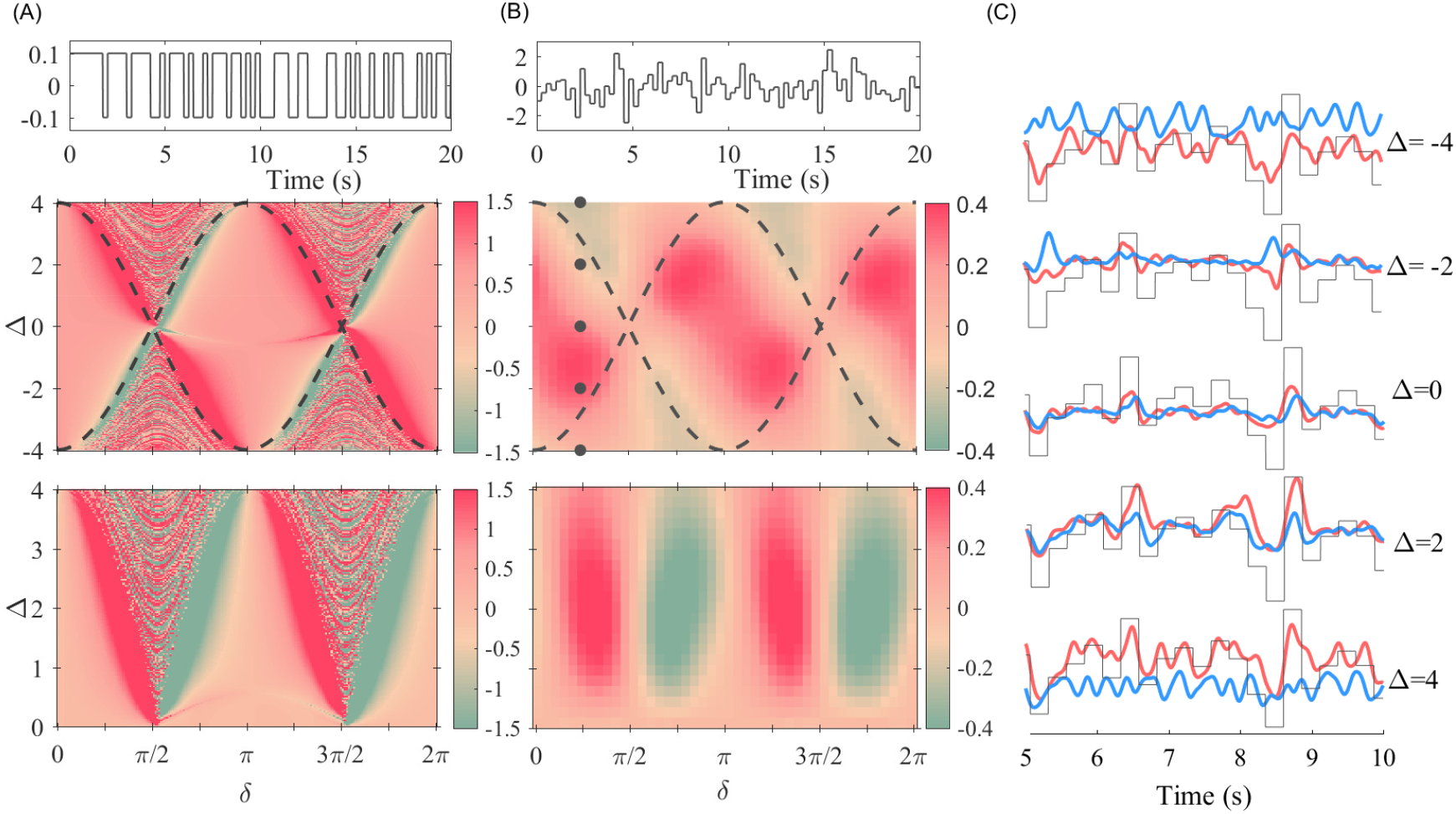
Numerical results for the case of Kuramoto oscillators. In panel (A) top we plot the stochastic dichotomous signal that is applied to the first oscillator (the sender). In panel (A) middle, we plot the correlation of the rate of the phase change of the second oscillator (the receiver) with the signal. The bottom panel is the transmission imbalance (eq. 7) which quantifies the difference of the signal transmission in the two directions. In panel (B) top we plot the random signal that is applied to the sender oscillator. In the middle panel, we plot the rate of the phase change of the receiver oscillator with the signal and bottom panel is the transmission imbalance. (C) Rate of phase change of two coupled oscillators for *δ* = *π*/4 and Δ = 0, ±2, ±4. The black line is the scaled input signal. The simulation parameters: *ω* = 55*Hz*, *K* = 4, *δ* ∈ [0, 2*π*], Δ = 4, and we added noisy input with *μ* = 0, and *σ* =1.

### Analysis

In our analysis, we calculated the firing rate (multi unit activity (MUA)) of the populations by using a Gaussian time window with *σ* = 2 *ms* and *σ* = 100 *ms* for fast and slow modulation, respectively. By moving in time the Gaussian time window, we calculated the weighted sum of the number of spikes in the window and put the value of it to the related time at the center of the time window.

#### Cross covariance

The cross-covariance quantifies the similarity between two vector. We used this measure to find the similarity between the firing rate of the receiver population with the input signal that we injected on the excitatory neurons of the sender (host) population. We assumed that if the signal was transmitted to the second population, its behavior (firing rate) should follow the signal. In the figures we plotted the un-biased and un-normalized value of the cross-covariance at zero lag (ZLC).

#### Coherency

To find the coherency of the populations we used a Gaussian time window with *σ* = 2(*ms*), to calculate the firing rate of the population activity using the spike train of the neurons. We defined the maximum coherency (*C* = 1), if all neurons spiked at the same time. We calculated and preserved the maximum value of the firing rate. We defined the coherency level of the oscillation by finding the firing rate of the population activity (during 2 sec) and taking average over the last 20 oscillations, normalizing to the value of the firing rate at the maximum coherency.

#### Delayed mutual information

To characterize the signal transmission and to define the effective connectivity between the coupled populations, we calculated the time delayed mutual information (dMI) [45], that quantifies the causal relation between the activities of the coupled populations. The time delayed mutual information is computed based on the Shannon’s entropy for two vector as:

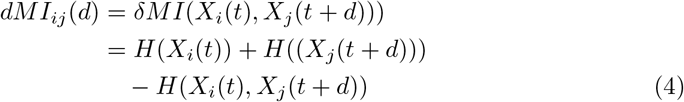

where 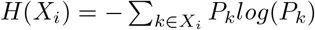 is the marginal entropy of *X_i_* and log is base 2 logarithm. *P_k_* is the probability of occurrence of event k. The Joint entropy in Eq. 4 i calculated as

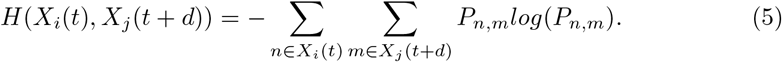

By integrating the dMI for positive lags, we quantified the amount of information that was transmitted from *i* to *j*; integrating over negative lags, we computed the transferred information in the opposite direction. Subtracting these two values, we obtained the resulting information transferred between the two populations.

#### Phase Response Curve (PRC) of a population

To find the response of a population to an injected pulse, we proceeded in a similar way as we do when we want to find the PRC of a single neuron. First we calculated the firing rate of population activity. Then we partitioned the time between two selected peaks into 30 segments. For each of these segments, and by keeping all parameters unchanged, we applied a rectangular pulse at a specified phase into all excitatory neurons of the population. By recalculating the firing rate of the population, we defined the PRC as the difference between the osillation period of the population without and with the injected pulse. The width and amplitude of the rectangular pulse that we used were 2 *ms* and 1 *μA/cm*^2^, respectively.

## Results

We start by investigating the role of the combination of transmission delay and frequency mismatch on the effective communication between two bidirectionally connected neural populations. Each population consists of N neurons (80% excitatory and 20% inhibitory) modeled by the Hodgkin-Huxley (HH) equations. The intra-population connectivity probability is 10% for all type of connections. Long-range excitatory projection connect the excitatory neurons of the two populations with a probability of 5%. The synaptic currents, assumed to be mediated by *AMPA* and *GABA_A_* receptors, are modeled by a double-exponential function with synaptic rise and decay time equal to 0.5 ms and 3 ms, respectively, for both type of synapses. The delay between any pair of connected neurons inside the populations is assumed to be 0.5 ms. The connection delay between populations will be varied. All neurons receive a Gaussian white noise with mean *μ* and variance *σ* (see Methods). In Fig. 1A-B, the coherence and the oscillation frequency of the isolated populations is shown, when changing the mean and the variance of the external noise. For the set of parameters we have chosen (dots in Fig. 1A-B and Table 1), each population exhibits synchronous oscillations in the high-gamma range. Sample spiking activity of the neurons and the population activity of the networks are shown in Fig. 1C.

### Phase locking between populations

We now concentrate on two populations connected with a given delay time (Fig. 1A). The mean input current into population 2 is fixed at 11*μA*/*cm*^2^ (unless otherwise noted), while the mean input current into population 1 is set at 11 ± Δ, where Δ introduces a frequency mismatch (detuning) between the two populations. We hypothesize that the theory of coupled oscillators can qualitatively predict the properties of a system of two populations connected via long range projections [6, 26, 30, 39, 42–44]. Then the degree of coherence between local oscillations of the two populations is determined by the coupling strength and transmission delay of the connections as well as the mismatch between the oscillation frequencies of the populations [20, 29]. In this study we fix the strength of the long-range connections and vary the frequency mismatch and the time delay. We observe that the locking window, determined by the maximum frequency mismatch for which the system remains in the phase-locked regime, depends on the delay, as expected (Fig. 2B-left panel). Within the locking zone, where the two coupled populations oscillate at the same frequency, while the phase difference between their oscillations changes with the delay and the frequency mismatch (Fig. 2B-right panel). Such a varying phase difference affects the signal and information transmission between the two populations as is shown below.

### Transmission of slow (rate) signals

To evaluate the ability of the system to transmit information, we apply a time dependent signal on one of the populations (the sender) and check how this signal is transmitted to the other population (the receiver). We consider the two widespread neuronal coding schemes, rate and spike-time coding. We do this by applying two different types of signals. In the first case we modulate the oscillation frequency of the sender population by using a time dependent input current (only on the excitatory neurons) whose frequency is much lower than the oscillation frequency of the populations (Fig. 2C-E).

To assess the quality of the signal transmission, we first extract the instantaneous oscillation frequency of the two populations, and then calculate the cross covariance between the rate of the receiver population and the signal (see Methods). The result is shown in Fig. 2C. Red color indicates a good transmission quality while green denotes that the transmission is degraded. Some aspects are highlighted in Fig. 2E. A dominant red color is observed for positive detuning indicating that the signal is better transmitted when the sender oscillates at a higher frequency than the receiver for most values of the transmission delay. However, also areas of weak transmission can be also seen for some specific delay values and positive detuning. For negative detuning, signal transmission can also occur from the population with lower oscillation frequency to that of high oscillation frequency for certain delays with a relatively good, although not maximal, quality. There are also some delay values that permit transmission in both directions with a relatively good transmission quality. Finally, the zero detuning case is not an optimal choice to transmit the signal. Similar results are obtained when computing the delay mutual information as shown in Fig. 2D (see below).

Time traces of the evolution of the populations’ rate superimposed on the signal, are shown in Fig. 2E for different delays and frequency mismatches (black dots in Fig. 2C). It is seen that for small delays the signal transmits better from the population that oscillates at higher frequency to the one that oscillates at lower frequency, as reported in previous studies (Fig. 2C and the left column of E) [19, 24, 40]. However, this no longer holds for larger delays. Indeed, we observe that, for some values of the delay, the quality of the transmission can be almost the same for both positive or negative detuning (Fig. 2C and middle column of E) or can be even better for negative detuning, i.e., when the sender population oscillates at a lower frequency (Fig. 2C and the right column of E).

### Transmission of pulse packets

In the second case we apply a single pulse packet on all excitatory neurons of the sender population at a certain phase (defined between 0 and 2*π* over one cycle of the oscillation) and measure the change in the response in both the sender and receiver populations. The phase at which the pulse is applied is varied in order to cover the whole 2*π* range. The effect of these pulses in the sender population is characterized by the local phase response curve (pPRC) while that in the receiver population is quantified by the non-local phase response curve (nPRC) which the latter is a measure for signal transmission.

In the different panels of Fig. 3A the pPRC and the nPRC upon the impact of pulse packets are shown, for different values of delay and frequency detuning. It can be clearly seen that for certain delay values, the nPRC has a finite amplitude which indicates the transmission of the signal. However, for other delays, the nPRC is flat indicating that the signal has not affected the receiver population. We also integrated the absolute value of the nPRC curves over a period as a measure of the signal transmission quality (results shown in Fig. 3B). It can be seen that the results qualitatively agree with those obtained for slowly varying signals (rate modulation; see Fig. 2). For small delays the signals are more efficiently transmitted from the fast to the slow oscillating population, but for larger delay values, symmetric transmission or even a better transmission in the reverse direction are found. It should be noted that, as occurs in the case of slow modulation, a better signal transmission is found in general for positive detuning (higher oscillation frequency of the sender population) as compared to the case of negative detuning (lower oscillation frequency of the sender population) when changing the coupling delay.

Green curves in In Fig. 3A and Fig. 3B show the prediction of the analytical results based on the multiplication the pPRC and the absolute value of its derivative, at the time at which the spikes of the sender populations arrive to the receiver population (after the delay, see section Theoretical background). A good agreement is seen between the analytical predictions and the numerical results.

### The PRC qualitatively predicts the information transmission flow

Our previous results highlight the importance of the PRC analysis. Indeed, the response of the receiver population to a perturbation applied to the sender population depends on the excitability state of the both populations at the time they receive the perturbation. The value of the PRC at the phase at which the external pulse impacts the sender populations determines the local response and the effect of the pulse on the sender population. If the receiver population receives the perturbation (after a transmission time δ) in a phase at which the time-derivative of the PRC is non-negligible (within the non-zero part of the blue solid curves in Fig. 3A), then the receiver population detects the perturbation. Otherwise, the perturbation is filtered out. In Fig. 4A we have shown examples for how a perturbation is transmitted or filtered out by analyzing the PRC of the populations. In Fig. 4B we quantify the transmission quality by changing detuning and delay solely based on the PRC of the oscillating populations without any external perturbation (See also the section Theoretical background). It can be seen that the results agree well with those obtained by analyzing the pulse transmission in Fig. 3B, highlighting the importance of PRC analysis.

### Information transfer

In the previous section we used cross-covariance and phase response curves as measures of the quality of the transmission of external signals in the system. Here we question if the above results can predict (and be inferred from) the causal relationship between the populations and the direction of the information flow. To this end, we compute the delayed mutual information (see Methods Eq. 4) between the two populations. This measure quantifies the information flow regardless of how the information is encoded and decoded [45].

Previous studies have shown that, in the absence of transmission delay, a frequency mismatch between the oscillations of the two populations breaks the symmetry of the information flow favoring the fast-to-slow direction [19, 24, 40]. As it can be seen in Fig.2D the direction of information flow changes with the delay and frequency mismatch in a qualitatively similar manner as for the signal transmission, indicating that the quality of signal transmission can accurately predict the direction of information flow and vice versa.

It is worth mentioning that similar results can be obtained when the networks oscillate in another frequency, being the only difference that the range of delays scales with the carrier frequency. This is a remarkable result due to its functional importance: When taking into account the transmission delays, the quality of signal transmission depends on the (carrier) frequency.

### Effect of asymmetric connectivity

The connections between brain regions are mostly asymmetric [46, 47]. Due to its biological importance, we check the results for the case of asymmetric connection strengths between two populations. We first explore the results for a feedforward network (no feedback connection) from the receiver to the sender. The computed mutual information transfer reveals that the transmission is independent of the delay, as expected, and is determined by the mismatch between the oscillation frequencies of the two populations (Fig. 5A). For all delay values, the positive mismatch (higher frequency of the sender population) yields a better information transfer.

We then fix the detuning at a positive value and the connection from sender and receiver to *g_for_*, and vary the strength of the connection from the receiver to the sender from zero to *2g_for_*. The effect of the delay becomes more apparent with increasing feedback strength (Fig. 5B). For *g_bac_* < 0.5*g_for_* the information transmission remains from the sender to the receiver almost independent of the delay. For *g_bac_* > 0.5*g_for_* we find windows of delays where the information transfer from the sender to the receiver is considerably degraded, while in other ranges the transmission is facilitated. Interestingly, the presence of the feedback connections facilitate the transmission for some ranges of delay and degrades the transmission for some other ranges.

### Theoretical background

To gain insight into the mechanisms that regulate the information flow between to delay-coupled oscillating neuronal populations, we analyze a minimal model of two coupled phase oscillators. These oscillators are characterized by their natural frequency *ω_i_* and their phase response function *Q_i_*. The evolution of the system is described by:

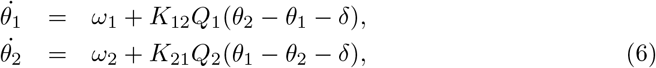

where *θ*_1_ and *θ*_2_ are the phase of the oscillators, *K*_12_ and *K*_21_ are the coupling strengths, and *δ* represents the interaction phase (that relates to the delay *τ* as *δ* = *ω_locked_* * *τ*). We assume that the strength of the connections are symmetric K_12_ = *K*_21_ = *K* and that response functions are the same *Q*_1_ = *Q*_2_ = *Q*. We then define the new variables *ϕ* = *θ*_1_ – *θ*_2_ and Θ = *θ*_1_ + *θ*_2_ and find

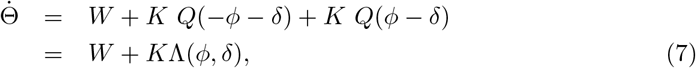

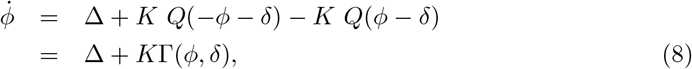

where, Ω = *ω*_1_ + *ω*_2_ and Δ = *ω*_1_ – *ω*_2_. Phase-locking is then determined by *ϕ* = 0. The phase difference in the locked state is implicitly given by

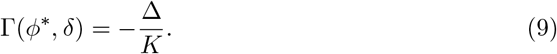

A solution of this system exists while 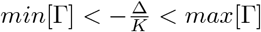, and the stability condition for a locked state is given by 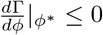.

The main objective is to explore how a *local* external signal imposed on one of the oscillators affects the other, i.e., how the signal *transmits*. The signal appears as a weak time dependent perturbation on the intrinsic frequency of one of the oscillators-the sender oscillator. Now we address the question of how the signal injected into one oscillator (the sender) affects the other oscillator (the receiver). Here we consider both the tonic signals which vary in a long time scale compared to the period of the oscillations, and pulsatile signals that model the synaptic inputs whose time constants (mainly decay time) are short compared to the oscillation periods of the populations. These approaches can be related to the rate and spike-time coding schemes in neuroscience [48].

To quantify the transmission of synchronous signals, we define a non-local phase response curve (nPRC) which is specified as the change in the phase of the receiver oscillator upon the incidence of the pulse injected into the sender one [49]. We assume that the unperturbed oscillators are locked at the phase difference *ϕ*^*^(Δ, *δ*) determined by Eq. 9. The impact of a pulse at a given phase *β* on the sender oscillator, changes its phase as Δ*ϕ*_1_ = *Q*(*β*). This gives rise to an instantaneous change in the argument of the coupling function in the second equation of Eq. 6 by *Q*(*β*) and changes the right hand side of that equation by *Q*(*β*)*Q′*(*ϕ*^*^ – *δ*) (where *Q′* is derivative of *Q* with respect to its argument), given *Q*(*β*) is small. As a result, the phase change in the receiver oscillator (and the nPRC) is expected to be proportional to both *Q*(*β*), which quantifies how much the sender is affected by the signal, and to *Q′*(*ϕ*^*^ – *δ*), which quantifies to what extent a change in the phase of the sender is transferred to the receiver (see Fig. 3A and B).

In the second case we consider a slowly varying signal and quantify the transmission of the signal by calculating a non-local response function defined as the derivative of the rate of the collective phase change Θ with respect to the free-running frequency of the sender oscillator *i*

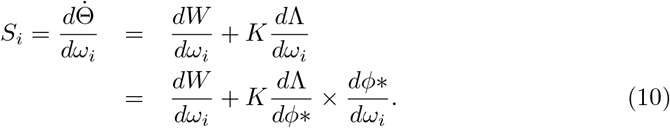

Note that the signal is assumed to be weak enough so that the system remains in the locked state, therefore, the dynamics of the collective phase is also representative of the dynamics of the receiver oscillator. The response function can be considered as a measure of the impact of the signal on the dynamics of the receiver oscillator. We will show through numerical simulation of Eq. 6 that this quantity can indeed qualitatively predict the correlation between the signal and the rate of change in the phase of the receiver oscillator.

As an example we consider *Q*(*α*) = *sin*(*α*) which serves as the canonical form for type-II excitable systems and resembles the interaction function for Kuramoto Model [50]. In this case, the phase difference *ϕ*^*^ in the locked state is determined by sin(*ϕ*^*^) = Δ/2*K* cos(*δ*) (Eq. 9), provided that |*W*/2*K* cos(*δ*)| ≤ 1. The non-local PRC, which quantifies the transmission of pulse signals, is proportional to cos(*ϕ*^*^ – *δ*). For the slow (rate) signals the response function is given by:

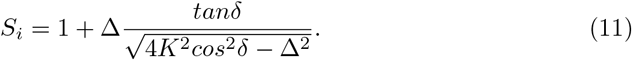

The corresponding imbalance measure is calculated as:

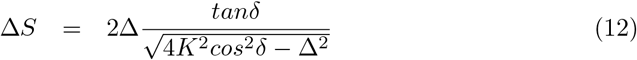

The analytical results for the nPRC and the response function *S_i_*, for different values of frequency mismatch Δ and delay *δ*, are plotted in Fig. 6A and B.

The results show that in the locked state, and for relatively small delays 0 < δ < *π*/2, the signal is better transmitted when it is injected on the high-frequency oscillator. Instead, for larger delays, phases *π*/2 < *δ* < *π*, the transmission is facilitated in the reverse direction from the low- to the high-frequency oscillator. Interestingly, the maximum response (and the minimum response in reverse direction) is found near the boundaries of locking zone, where the imbalance is also maximized.

To check the validity of our analytical results, we perform numerical simulations for the coupled phase oscillators with *Q*(*ϕ*) = *sin*(*ϕ*). In the first case we impose an impulse on the sender oscillator and numerically measure the phase change in the receiver oscillator. The results shown in Fig. 7 are in agreement with the theoretical prediction Fig. 6. We then apply a small amplitude dichotomous random signal, which switches between two states at random times, on the sender and calculate the correlation of the rate of the oscillations of the receiver with the signal. The results shown in Fig. 7(A) are in excellent agreement with the analytical results. The only difference with the analytical results is that for the out of phase solution, and near the locking region, a reliable transmission and a large imbalance is seen. However, since the analytical results are obtained assuming phase locking, they are not valid for this region. Consequently, the extension of the results to this region can be explained by the presence of intermittent locked epochs when the system approaches the locking zone [51].

Although the results normally depend on the exact form of PRC, a qualitative agreement with the simulation results for two neural populations (Fig. 2) can be observed since the pPRC of the population is type-II. But since the two PRCs are not exactly the same, the results do not conform in details, e.g., the transmission is not symmetric in the two directions over different delay ranges and favors fast-to-slow direction (compare Fig. 7 with (Fig. 2)).

## Discussion

Extensive experimental and theoretical studies over the recent decades have unleashed the role of brain oscillations in several cognitive and executive brain functions like sensory processing [52], memory [53, 54] attention [55], and motor functions [56]. Oscillations and the coordinated activity of the neurons facilitate the transmission of signals along different stages of neural processing systems and enable an efficient communication between brain areas [57–59]. Integration of information which is processed across distributed specialized brain regions is hypothesized to be controlled by the temporal coordination of their local dynamics [60]. Oscillations change the excitability of the neurons over time and enable control of communication between brain areas by adjusting their phase relations [18]. Moreover, oscillations provide a functional substrate to transmit multiplex of information along different routes and directions over different frequency bands [61–64].

Numerous theoretical and numerical studies have been carried out to shed light on the mechanisms through which the oscillations control the transmission of information and give rise to flexible communication channels in brain circuits [19, 20, 58, 65]. In the last years, the effect of the frequency mismatch, in particular, has been the focus of several theoretical studies addressing the control of information routing in brain circuits [19, 20, 24, 66, 67].

In this paper, we revealed the essential role that the transmission delay plays in the efficiency and direction of functional interaction between the oscillating regions. With a systematic change of the frequency mismatch and delay time in the coupling between two oscillating neuronal populations, we showed that previous results on the information flow from high-frequency to low-frequency populations is only correct for small delays [20, 24]. Interestingly, other patterns of effective communication, including almost symmetric communication channel or information flow in the reverse direction, can be observed when increasing the delay. Notably, our results were found to be valid and consistent for both slow (rate coding type) and fast (temporal coding type) signals.

### 0.1 Role of collective phase response functions

The PRC has been extensively shown to be important to explain different synchronization scenarios between coupled dynamical systems (see .e.g. [29, 68]). It the case of delay-coupled neuronal populations, the PRC accurately reveals the efficacy and the direction of the information transfer. Our theoretical framework with phase oscillators showed that the signal transmission depends on the phase response of the coupled elements (phase oscillators or neural populations). Once knowing the collective PRC of the neural population (pPRC), the regions for good transmission of the signals in the parameter space including delay and the frequency detuning can be predicted. These results were in good agreement with those obtained when modulating the sender population and computing the information flow.

While the collective phase response is widely studied for populations of oscillators [69–72], those results are not readily applicable to the neural ensembles. The neural populations composed of excitatory and inhibitory neurons show collective oscillations at the population level while the single neurons irregularly fire [11, 73].

The mechanism of the synchrony and mathematical formulations of the emergence of the rhythms in such networks are far different from those of networks of coupled oscillators [74, 75]; though their response to the external inputs might be different and warrants for more systematic studies [49, 76].

### 0.2 Role of the frequency detuning and connection delay

While the role of frequency mismatch has been shown to be an important factor in information routing [19, 20, 24, 66, 67], here we showed that, more than the detuning alone, is the combination between the detuning and the connection delay which matters. Based on our results, it can be concluded that there is a certain preference towards a larger flow of information for positive detuning than for negative ones compatible with previous studies. But this is not the case for all values of the delays. Symmetric information exchange and information flow from the low to the high frequency population is also possible in some narrower range of delays.

### 0.3 Limitations and future studies

In this study our network operated at the state of oscillation at a single frequency. We showed that the signals and information transmission change with delay, when delay is changed over a period of oscillation. At a given delay time, then, the transmission changes with frequency of oscillation. It can be concluded that the effective connectivity between the brain regions is different at different frequencies because of the delayed interactions. This provides the possibility to transmit the information over different routes and directions at different frequency bands as is observed in experimental studies [61, 62]. Since our network was capable to produce oscillation in single frequency in the gamma range, it was not possible to check the transmission in multiple frequency bands.

In the brain, the network operates at multiple frequency bands (including a fast and a slow oscillatory component) and several modeling studies have suggested mechanisms to reproduce this regime [77–79]. In our model we only considered synapses mediated by *AMPA* and *GABA_A_* receptors, and used the classical description of the Hodgkin-Huxley neuron model. To account for networks with richer dynamics and to produce oscillations at multiple frequencies, we need to incorporate synapses with slow dynamics and chose more appropriate neuronal models [77, 80]. Moreover, to highlight the role of delay we set our network in a high coherence regime. It is necessary to explore how the results translate to more realistic networks with unstable oscillatory dynamics and lower coherence [19].

## Acknowledgments

This work was partially supported by the Spanish State Research Agency, through the Severo Ochoa and María de Maeztu Program for Centers and Units of Excellence in R&D (MDM-2017-0711), MINECO (Spain) through project TEC2016-80063-C3 (AEI/FEDER, UE) and Ministerio de Ciencia e Innovación (AEI/FEDER, UE) through projects PID2019-111537GB-C21 and PID2019-111537GB-C22. A.V. acknowldges partial support from Iranian Cognitive Sciences and Technologies Council, under the grant No. 832.

